# Multivariate Analysis Of Potato Genotypes For Genetic Diversity

**DOI:** 10.1101/2020.04.14.041236

**Authors:** Arifa Khan, Shazia Erum, Naveeda Riaz, Muhammad Ibrar Shinwari, Maryum Ibrar Shinwari

## Abstract

Sustainable production of food crops relies on germplasm improvement and genetic diversity that helps to identify appropriate parents, which is very important step in breeding of genotypes having high yield potential for future use. This study was conducted to investigate the extent of genetic diversity using multivariate technique on the basis of qualitative and quantitative traits. An experiment was comprised of 74 exotic genotypes and started at National Agriculture Research Center Islamabad, Pakistan during autumn 2017-2018 and 2018-2019. Data was recorded on qualitative and quantitative traits by following standard procedures and Biplot analysis was used to calculate the significance among the studied quantitative traits to exhibit the strength of relationship between traits. Results showed significant diversity in qualitative traits and quantitative traits. Red, yellowish, brown, light yellow, light brown color tubers were produced. Alike, genotypes produced yellow, cream and white flesh color tubers. Genotypes had oval, round, oblong, elliptic and reform with medium, small and large size tubers. Alike, brown, light brown, dark red and yellow eyes color was noted. In case of quantitative traits, genotypes had high variance regarding plant height, leaf area and number of tubers per lane. Genotypes had very high genetic variance for weight of tuber per plant and weight of tuber per lane while low variance was recorded for germination, number of stem per plant and number of eyes per tuber. Significant positive correlation was observed between number of tubers per plant (TPP) with number of eyes on tubers (r = 0.241) and number of tubers per lane (TPL) (r = 0.349). But negative correlation was noted between number of tubers per plant (TPP) with plant height (r = - 246), leaf area (−0.529) and germination (r = −0.283). Plant height was found significantly positive correlated with leaf area (r= 0.456), germination percentage (r = 0.255) and weight of tubers per plant (r = 0.307). Leaf area (LA) showed positive significant correlation with number of tubers per plant (r = 0.466) and weight of tubers per plant (r = 0.263)., yield and harvest index (r = 0.798, 0.755, 0.255). Weight of tubers per lane (WTL) showed positive correlation with weight of tubers per plant (r = 0.387). Regarding the interrelation between the traits and genotypes, the first two principal component axes (PC1, 24.83% and PC2, 23.46%) accounted for about 48.29% of the total variability reflecting the complexity of the variation between the plotted traits of genotypes.

## Introduction

The availability of diverse genetic germplasm ensures success in development of new high yielded cultivars (Buckler, 2009). Genetic diversity and suitable germplasm are very important factor in agricultural crops. Genetic diversity helps to identify appropriate parents, which is very important step in breeding of genotypes having high yield potential for future use. A comprehensive knowledge about variability in genetic material is necessary for the improvement of suitable traits (Kahrizi, Maniee, Mohammadi, & Cheghamirza.., 2010). Knowing and understanding the accessible diverse genetic germplasm is of the most important for successful exploitation and estimation of germplasm (Zubair, Ajmal, Anwar, & Haqqani, 2007). The genetic diversity is helpful to identify different parental combinations to produce progenies having maximum genetic variability for further selection (Mohammadi & Prasanna, 2003) and one can take genes of our choice from diverse germplasm (Habtamu, 2013). Genetic relationships between pure or inbred lines is also important for scheduling crosses, selection of lines to particular heterotic groups and for precise recognition regarding plant varietal protection (Franco et al., 2001). Genetic diversity also assists to identify such groups possessing similar genetic background and their utilization as a genetic resource (Thompson, Nelson, & Vodkin, 1998).

Multivariate analysis is an important and very popular method for estimation of genetic variability(Malik et al., 2014) and to determine pattern of dissimilarities and their genetic relationship between germplasm collections (Ajmal et al., 2013). Multivariate analyses have been used in various countries (Babić, Pajić, Prodanović, Babić, & Filipović, 2010) for different food crops like wheat (Ajmal et al., 2013), maize (Azad, Biswas, Alam, & Alam, 2012; Lee, Herrman, Lingenfelser, & Jackson, 2005) and sorghum (Ali et al., 2011). To select genetically distance parents, various genetic diversity researches have been initiated between crop species on the basis of quantitative and qualitative traits (Hailegiorgis, Mesfin, & Genet, 2011). To harness friable genetic variation in breeding material, it is worthwhile to trace the total variation into its components. The present research was initiated realizing the significance and need for such a comparative study in potato particularly to investigate the level of genetic diversity by employing multivariate technique on the basis of qualitative and quantitative characters to sort out superior genotypes and to adopt a suitable breeding program for variety development in country. Keeping this in view, the present study was carried out to evaluate their genetic diversity.

## Materials and Methods

The experiment was evaluated at Plant Genetic Resource Institute (PGRI), National Agriculture Research Centre (NARC) Islamabad, Pakistan during November 2017-2018 to 2018-2019. In this experiment, 76 exotic genotypes (Table 1) were imported from International Potato Centre, Peru. The experiment was sown in a RCBD (Randomized Complete Block Design) with plant to plant and row to row distance of 25cm and 65 cm, respectively. The recommended dose of fertilizers i.e. nitrogen 250 kg/ha, phosphorus 125 kg/ha and potassium 125 kg ha^-1^ was applied. All the phosphorus, potassium and half dose of nitrogen were applied at the time of sowing while remaining was used at 1st and 2^nd^earthing up. Crop was visited regularly during growing season. Irrigation and plant protection measures were carried out when required. Observations for qualitative traits such as tuber shape, tuber size tuber color, tuber flesh color, tuber skin, eye color of tuber and tuber eyes depth were taken. Similarly, quantitative traits such sprouting percentage, plant height, number of stem, leaf area, number of tubers/plant, number of tuber/row, weight of tuber/plant, weight of tuber/row and number of eyes/tuber were recorded by following standard procedures.

**Table 1.** Genotypes studied during experiment

| Sr. No. | Genotypes | Sr. No. | Genotypes | Sr. No. | Genotypes | Sr. No | Genotypes |
| --- | --- | --- | --- | --- | --- | --- | --- |
| 1 | CIP1 | 20 | CIP20 | 39 | Frolica | 58 | G-128 |
| 2 | CIP2 | 21 | CIP22 | 40 | Miss Andes | 59 | N-U-II-1 |
| 3 | CIP3 | 22 | CIP24 | 41 | SL-8-5 | 60 | Torage |
| 4 | CIP4 | 23 | CIP25 | 42 | PRS-493 | 61 | Safari |
| 5 | CIP5 | 24 | CIP27 | 43 | Hi brid 25-1 | 62 | NU-II-J |
| 6 | CIP6 | 25 | CIP28 | 44 | RMS-5-81-43-R | 63 | NU-II-5 |
| 7 | CIP7 | 26 | CIP29 | 45 | N-18 | 64 | Zina RED |
| 8 | CIP8 | 27 | CIP30 | 46 | NU-IIF | 65 | HZD2 1499 |
| 9 | CIP9 | 28 | CIP31 | 47 | NUYT-Q | 66 | Asterix |
| 10 | CIP10 | 29 | CIP 32 | 48 | NU-110 | 67 | Desiree |
| 11 | C1P11 | 30 | CIP34 | 49 | NUYT-F | 68 | Karuda |
| 12 | CIP12 | 31 | Focus | 50 | NUIIB | 69 | Harmes |
| 13 | CIP13 | 32 | Missmagnone | 51 | NUIIC | 70 | Triplo |
| 14 | CIP14 | 33 | M.05 81-43R | 52 | NU-IIM | 71 | Melanto |
| 15 | CIP15 | 34 | Elbida | 53 | NU-IIA | 72 | Flamba |
| 16 | CIP16 | 35 | Nazka | 54 | NUYTL | 73 | Aestrix |
| 17 | CIP17 | 36 | Red Valentine | 55 | Daseer | 74 | Amarin |
| 18 | CIP18 | 37 | Sante | 56 | NIIH | 75 | Kuroda |
| 19 | CIP19 | 38 | Tourage | 57 | RRS-1035 | 76 | Stemster |

### Observations

Following observations were recorded during experimental trials.

For tuber shape, tuber size tuber color, tuber flesh color, tuber skin, eye color of tuber and tuber eyes depth, tubers of 5 plants of each genotype were taken and observed visually.

### Sprouting percentage

It was calculated by following formula

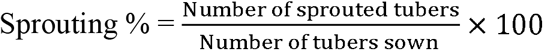

### Plant height (cm)

For plant height, 5 plants of each genotype were selected randomly and their height was measured from base to top with the help of a meter rod and then averaged.

### Number of stem/plant

For number of stem/plant, 5 plants of each genotype were selected randomly and their number od stem was noted.

### Number of tubers per plant and number of tubers per row

To count the number of tuber per plant, 5 plants from each genotype selected randomly and their tubers were counted and then averaged. At harvesting, all the potatoes of each genotype in their respective rows were counted.

### Tubers weight per plant and tubers weight per row (g)

For tuber weight, 5 plants of each genotype were selected randomly and their tubers weight was noted with the help of digital electrical balance and then averaged. At harvesting, all the potatoes of each genotype in their respective rows were weighted individually.

### Number of eyes

For number of eyes, tubers of 5 plants of each genotype were observed carefully after harvesting and their eyes were noted and then averaged.

### Leaf area (cm^2^)

For leaf area, leaves of 5 plants of each genotype were harvested, their leaves were separated and leaf area was noted with help of leaf area meter.

### Data analysis

For basic statistics, data were analyzed in Microsoft Office Excel 2010. To establish phenotypic similarity and dissimilarity, a Biplot analysis was carried out. Pearson’s correlation coefficient was also calculated and the significance was noted among the studied quantitative attributes to disclose the strength of relationship using XLSTAT 2012.

## Results and discussion

### Qualitative traits

Results showed that genotypes have diversity in qualitative traits (Figures 1–3). In case of color of tubers, maximum genotypes, CIP7, CIP28, M.05 81, Red valentine, SL-8-5, NUIIDPRS-493, Hi brid 25-1, RMS-5-81-43-R, NUYT-Q, NUIIB, NU-IIA, NIIH, RRS-1035, N-U-II-1, Asterix, Desiree, Karuda, had tubers of red color while CIP20, CIP22, CIP27, Focus, Elbida, Nazka, Sante, Tourage, Miss Andes, MISSMAGNONE and Frolica genotypes had yellowish color tubers(Figure 1). Genotypes CIP4, CIP5, CIP6, CIP13, NU-IIF, NU-IIM, NUYTL, DASEER, G-128, TORAGE, NU-II-J, NU-II-5 and ZINA RED had tubers of brown color. Light brown color tubers were harvested from CIP1, CIP9, CIP15, CIP16, CIP18, CIP19, CIP30, CIP31, CIP 32, NUIIC genotypes. CIP25 and N-18 genotypes produced light red color tubers. Similarly, majority of genotypes (CIP10, CIP13, CIP14, CIP15, CIP17, CIP18, CIP27, CIP29, CIP30, Elbida, Nazka, Sante, Tourage, Frolica, Miss andes, PRS-493, RMS-5-81-43-R, N-18, NU-IIA, NUYTL, NIIH, Asterix, Karuda, Focus and MISSMAGNONE produced tubers of yellow flesh color (Figure 1). While CIP2, CIP4, CIP6, CIP9, C1P11, CIP16, CIP22, CIP25, CIP28, CIP31, M.05 81, Red valentine, Hi brid 25-1, NU-110, NUYT-F, NUIIC, DASEER, N-U-II-1, TORAGE, MISS ANDES, NU-II-J, NU-II-5 and ZINA RED produced tubers with cream color flesh. Tubers with white color flesh were recorded for CIP1, CIP3, CIP5, CIP7, CIP19, CIP20, CIP 32, SL-8-5, NUIID, NU-IIF, NUYT-Q, NUIIB, NU-IIM, RRS-1035 and G-128(Figure 1).

**Figure 1.**
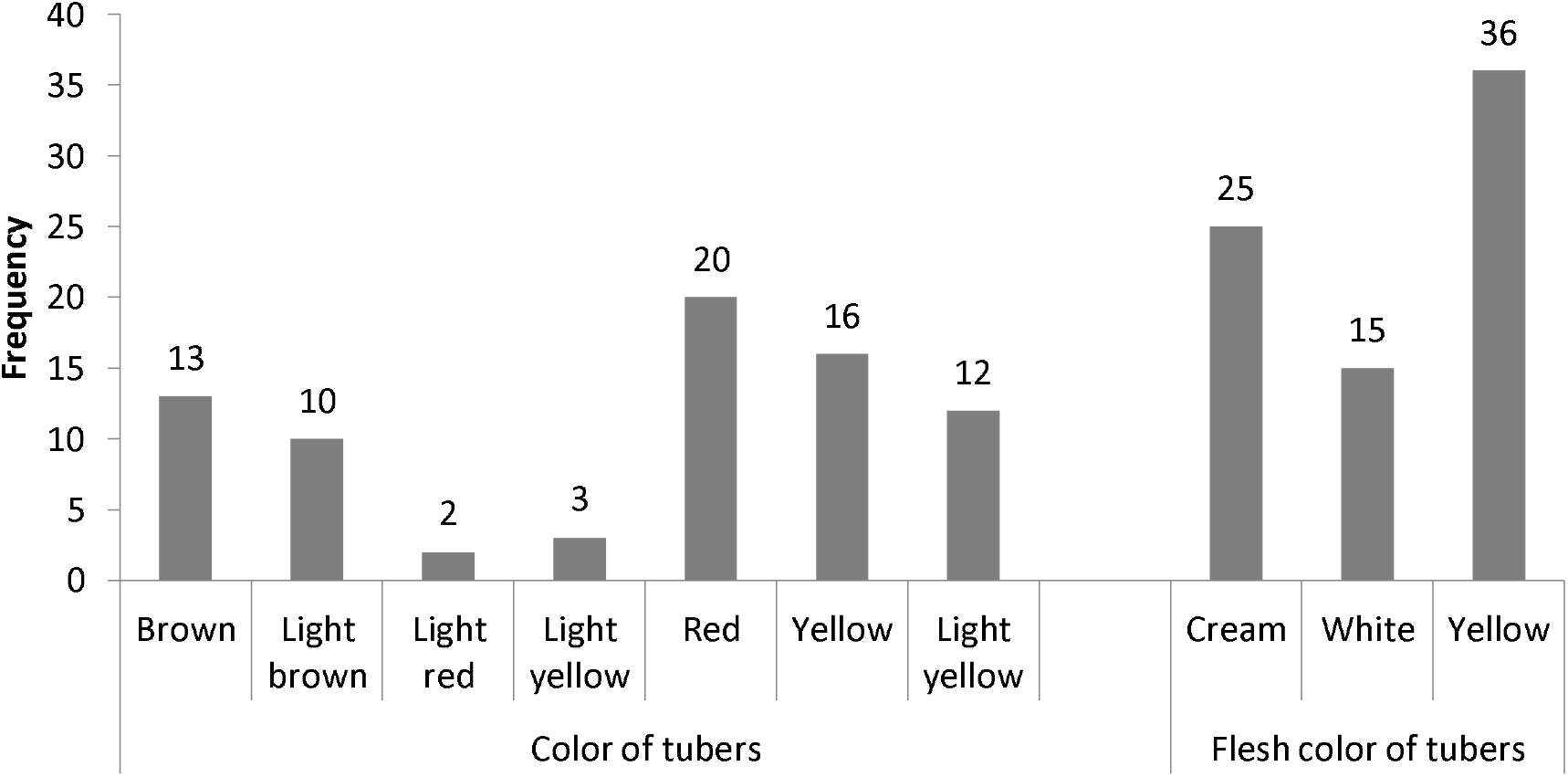
Tubers color and flesh color

**Figure 2.**
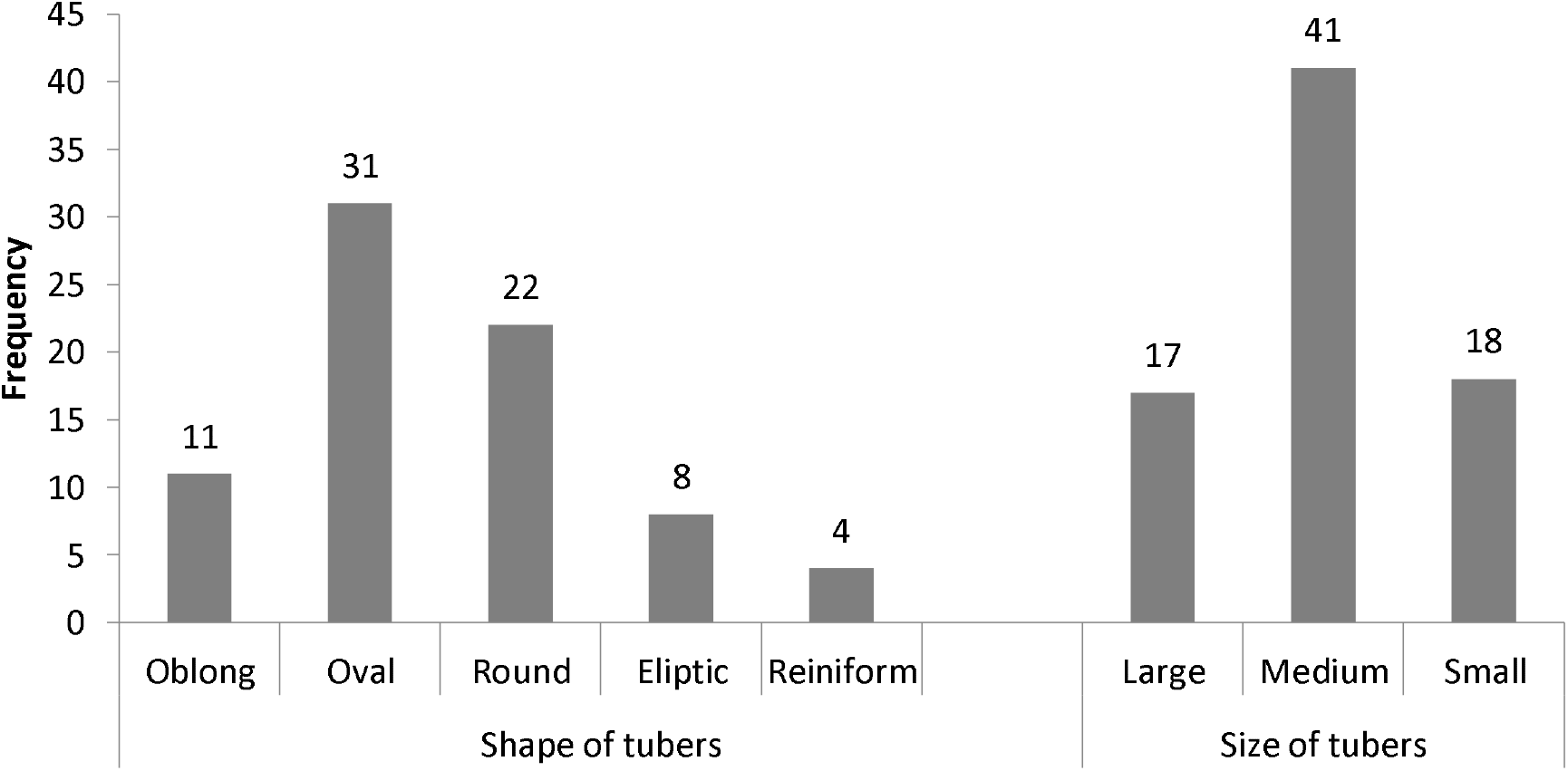
Size and shape of tubers

**Figure 3.**
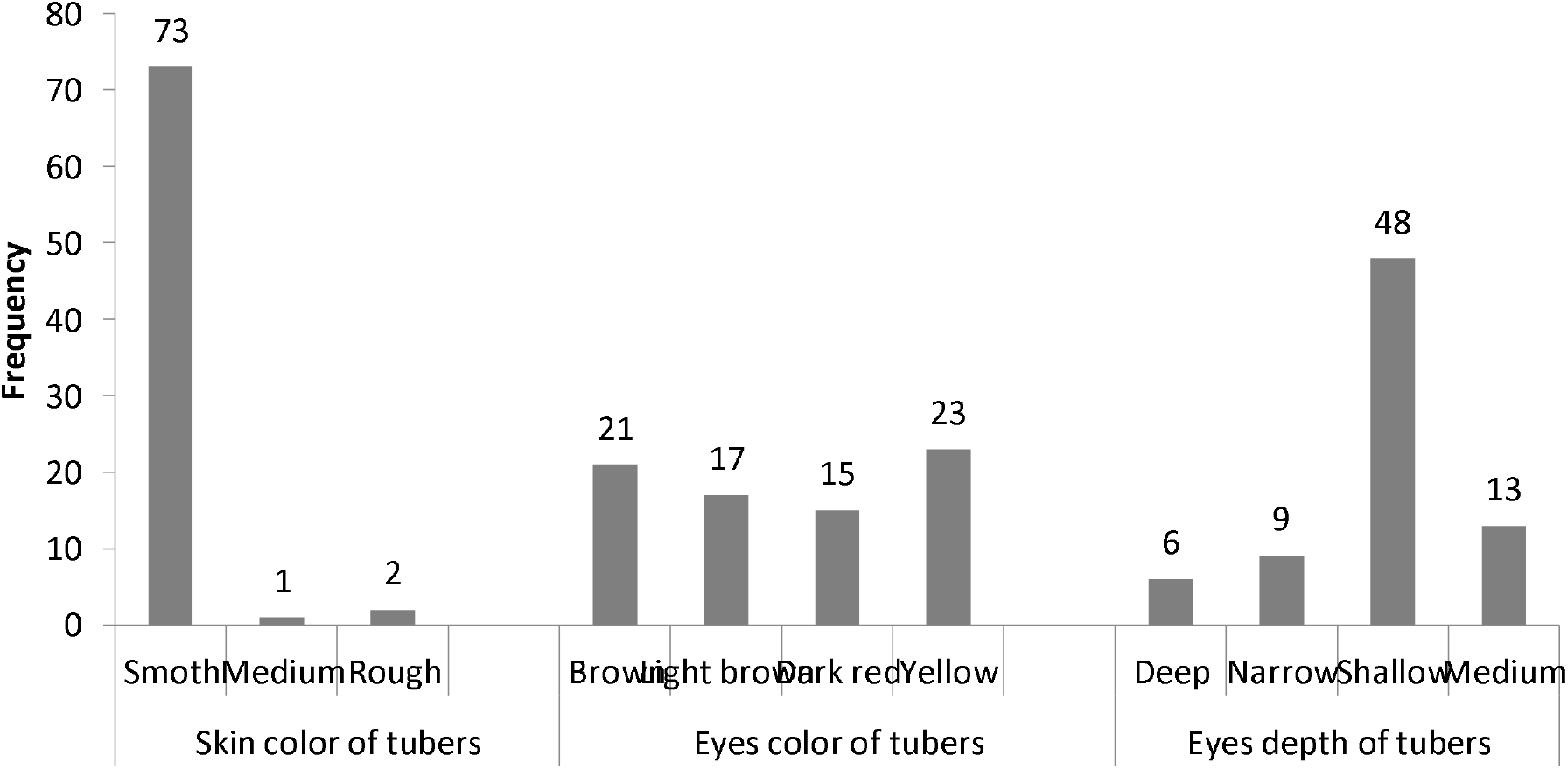
Skin, eye color of tubers and depth of eyes on tubers

In case of tuber shape, maximum genotypes (CIP2, CIP13, CIP14, CIP19, CIP25, CIP27, CIP34, Sante, SL-8-5, NUIID, PRS-493, RMS-5-81-43-R, N-18, NU-IIF, NU-110, NU-IIM, NU-IIA, RRS-1035, G-128, N-U-II-1, MISS ANDES, NU-II-J, NU-II-5, Asterix, Karudaand Desiree produced oval shaped tubers (Figure 2). While CIP1, CIP3, CIP4, CIP5, CIP6, CIP7, CIP10, CIP16, CIP17, CIP18, CIP20, CIP24, CIP28, CIP31, CIP 32, Elbida, Miss andes, Hi brid 25-1, NUYT-F, NUYTL and DASEER produced round shaped tubers. Focus MISSMAGNONE, M.05 81, Nazka, Red valentine, Tourage, Frolica and CIP29 genotypes produced elliptic shaped tubers. Alike, CIP3, CIP6, CIP8, CIP10, C1P11, CIP12, CIP13, CIP15, CIP16, CIP18, CIP20, CIP24, CIP28, CIP30, Focus, MISSMAGNONE, Nazka, Frolica, Miss andes, PRS-493, Hi brid 25-1, RMS-5-81-43-R, N-18, NU-IIF, NUYT-F, NUIIC, NU-IIM, NUYTL, RRS-1035, G-128, NU-II-J, NU-II-5and Asterix genotypes gave medium size tubers (Figure 2).CIP1, CIP2, CIP5, CIP7, CIP9, CIP14, CIP17, CIP22, CIP29, CIP31, Sante, SL-8-5, NUIID, NU-110, DASEER, N-U-II-1, TORAGE and MISS ANDES produced small size tubers while large tubers were noted for CIP4, CIP19, CIP25, CIP27, CIP 32, CIP34, M.05 81, Elbida, Red valentine, Tourage, NUYT-Q, NU-IIA and NIIH genotypes.

In case of skin color of tubers, all the genotypes except Asterix, CIP17 and CIP24 produced smooth skin tubers (Figure 3). In case of eye color, CIP13, CIP16, CIP20, CIP22, CIP29, CIP34, Focus, MISSMAGNONE, Elbida, Nazka, Tourage, Frolica, N-18, NUYT-F, NU-IIA, NUYTL, G-128, NU-II-5 and Karuda had yellow eye color tubers while light brown eye color tubers were recorded for CIP1, CIP9, CIP12, CIP14, CIP17, CIP24, CIP25, CIP31, Sante, RMS-5-81 −43-R, NU-IIF, NUIIB, Miss andes, CIP30, MISS ANDES and CIP4 genotypes. While, CIP2, CIP3, CIP7, CIP10, C1P11, CIP15, CIP19, CIP 32, PRS-493, RRS-1035, NU-II-J, Asterix and ZINA RED gave tubers with brown eye color. In depth of eyes of tubers, CIP5, CIP7, CIP10, C1P11, CIP13, CIP14, CIP16, CIP17, CIP18, CIP19, CIP22, CIP25, CIP27, CIP28, CIP31, CIP 32, Focus, MISSMAGNONE, M.05 81, Elbida, Nazka, Sante, Tourage, Frolica, Miss andes, PRS-493, RMS-5-81-43-R, NU-IIF, NU-110, NUIIC, NU-IIM, NUYTL, NIIH, RRS-1035, TORAGE, NU-II-5, Asterix, Desiree and Karuda had shallow eye depth. While, deep eyes were noted for tubers taken from CIP4, CIP34, Hibrid 25-1, N-18, NUYT-Q and NUYT-F genotypes (Figure 3).

### Quantitative traits

Seventy six potato genotypes (Table 1) were grown to evaluate the association or relationship among different parameters and to measure the diversity exiting in gene pool. Basic statistics for various traits are presented in Table 2. Results exhibited that genotypes had significant variation for quantitative traits. High variance was observed for plant height, leaf area of plants and number of tubers per row. A very high genetic variance was observed for weight of tuber per plant and weight of tuber per row while low variance was recorded for germination, number of stem per plant and number of eyes per tuber. Improvement of these traits through simple selection might be limited from genotypes used in the present study.

**Table 2.** Basic statistics for 9 quantitative traits of 76potato genotypes

| Traits | Maximum value |  | Minimum value |  | Mean |  | Standard deviation |  | Variance |  |
| --- | --- | --- | --- | --- | --- | --- | --- | --- | --- | --- |
|  | 2017 | 2018 | 2017 | 2018 | 2017 | 2018 | 2017 | 2018 | 2017 | 2018 |
| Germination % | 100 | 100 | 70 | 10 | 89.9 | 89.8 | 7.3 | 13.0 | 55 | 82 |
| Plant height | 84 | 86 | 22 | 23.6 | 48.8 | 50.7 | 14.3 | 14.7 | 216.8 | 219.2 |
| Leaf area/plant | 85 | 85.2 | 6 | 6.2 | 24.6 | 24.7 | 23.6 | 23.6 | 603 | 567 |
| Number of stem/plant | 11 | 11 | 2 | 2 | 5.00 | 4.9 | 1.6 | 1.60 | 6.1 | 2.4 |
| Number of Eyes/tuber | 8 | 8 | 2 | 2 | 5 | 5 | 1.3 | 1.24 | 3.5 | 1.6 |
| Tubers/plant | 23 | 23.4 | 4.4 | 4.4 | 10.9 | 10.5 | 4.4 | 4.63 | 28.9 | 21.7 |
| Tubers/row | 115 | 117 | 24 | 22 | 66.3 | 65.2 | 21.8 | 23.70 | 473 | 589 |
| Weight of tubers/plant | 783 | 1342.3 | 122.2 | 39.8 | 377.8 | 387.6 | 123.8 | 170.3 | 52241 | 32612 |
| Weight of tubers/row | 6841 | 6841 | 284.4 | 199.2 | 1563 | 1576 | 946.7 | 1021.4 | 9092967 | 935193 |

### Correlation analysis

Data regarding correlation is given in Table 3, which exhibited positive correlation between number of tubers per plant (TPP) with number of eyes on tubers (r = 0.241) and number of tubers per row (TPL) (r = 0.349). But negative correlation was noted between number of tubers per plant (TPP) with plant height (r = −246), leaf area (−0.529) and germination (r = - 0.283). Number of eyes present on tubers exhibited positive correlation with number of stem per plant (r = 0.282) but negatively correlated with germination percentage (r = −0.308) and weight of tubers per plant (r = −0.254). A positive correlation was recorded between plant height with leaf area (r= 0.456), germination percentage (r= 0.255) and weight of tubers per plant (r = 0.307). Leaf area (LA) depicted positive correlation with number of tubers per plant (r = 0.466) and weight of tubers per plant (r = 0.263)., yield and harvest index (r = 0.798, 0.755, 0.255). Weight of tubers per row (WTR) showed positive correlation with weight of tubers per plant (r = 0.387).

**Table 3.** Pearson’s correlation of genotypes for the estimated seven qualitative traits of wheat

| Variables | Eyes | Stem | PH | LA | TPR | WTR | Germ | WTP |
| --- | --- | --- | --- | --- | --- | --- | --- | --- |
| TPP | 0.241 | 0.166 | -0.246 | -0.529 | 0.349 | 0.304 | -0.283 | -0.092 |
| Eyes |  | 0.282 | 0.053 | 0.089 | 0.227 | -0.087 | -0.308 | -0.254 |
| Stem |  |  | -0.031 | 0.308 | 0.555 | 0.013 | -0.213 | 0.127 |
| PH |  |  |  | 0.456 | -0.030 | 0.179 | 0.315 | 0.307 |
| LA |  |  |  |  | 0.466 | -0.209 | 0.204 | 0.263 |
| TPR |  |  |  |  |  | -0.004 | -0.018 | 0.204 |
| WTR |  |  |  |  |  |  | 0.061 | 0.387 |
| Germ |  |  |  |  |  |  |  | 0.145 |

Values in bold are different from 0 with a significance level $\alpha=0.05$
TPP = number of tuber per plant, PH = plant height, LA = leaf area, TPR = number of tuber per row, WTR = weight of tubers per row, stem = number of stems per plant, Germ = germination percentage

### Biplot analysis

To identify the multivariate relationships and similarity in quantitative characters in 76 potato genotypes, biplot analysis was used (Figure 4). Regarding the interrelation between the genotypes and traits, the values of the first two PC axes (PC1, 24.83% and PC2, 23.46%) showed 48.29% of the total variability showing the diversity between the genotypes and plotted traits.

**Figure 4.**
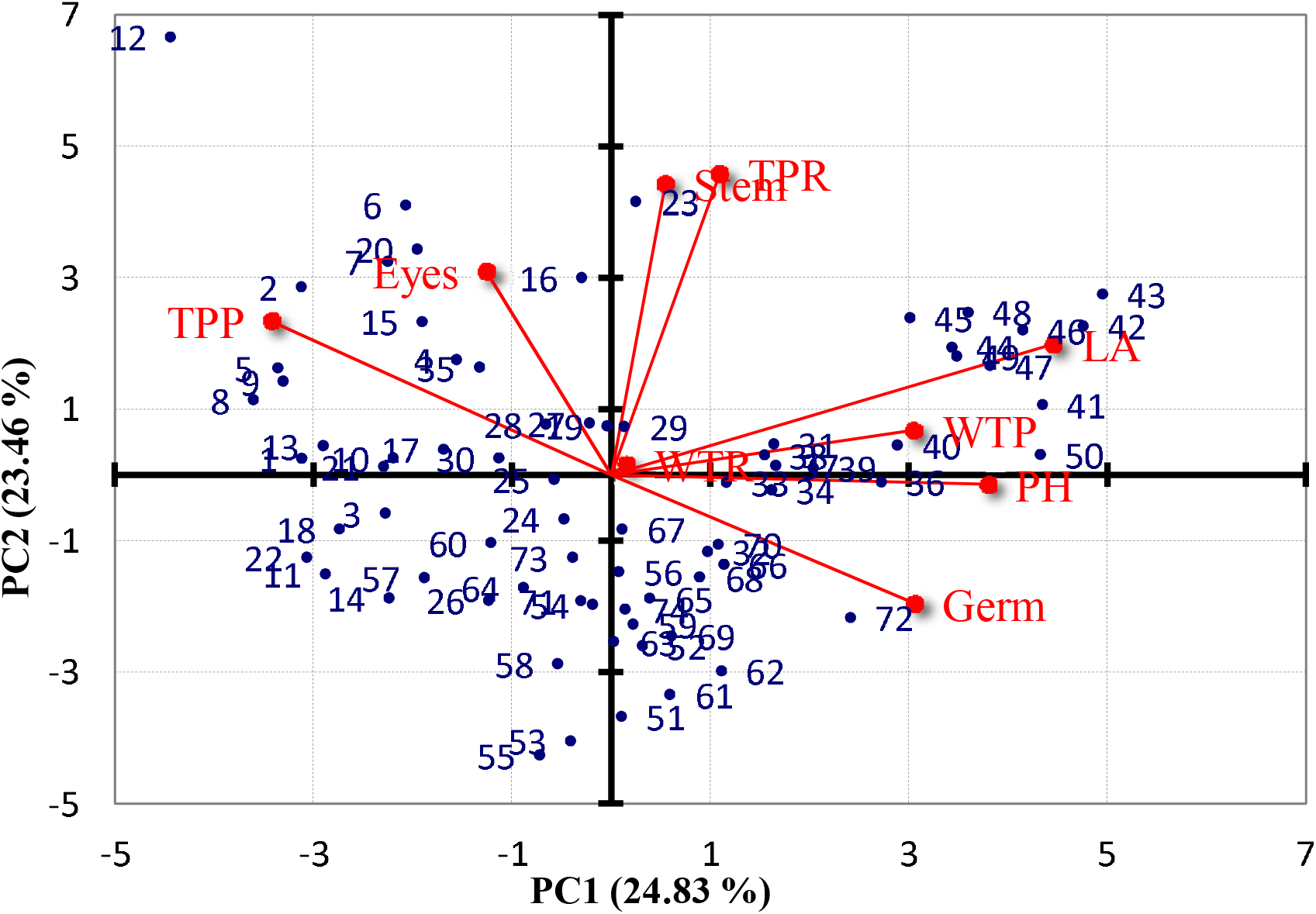
Biplot of studied traits and response of 76 potato varieties

Regarding the traits, PC1 had the number of stem per plant, (Stem), number of tubers per row (TPR), weight of tuber per row (WTR), leaf area (LA), weight of tuber plant (WTP), plant height (PH) and germination percentage (Germ) while, PC2 had the number of tubers per plant (TPP) and number of eyes on tubers (Eyes) as the primary elements. TRP and number of stems exhibited positive correlation in positive direction. Similarly, LA, WTP and WTL had high correlation in positive direction and TPR was also positively correlated with these traits in the positive direction while germination has correlation in the negative direction. Similarly, positive correlation was also observed between number of eyes and TPP. In contrast, number of eyes and TPP were negatively correlated with other quantitative parameters mainly LA, WTP, TPR and WTR.

Results showed that there is high variability among different genotypes such as yellowish, light yellow, red, brown, light brown regarding flesh color. Similarly, the size and shape of tubers was also different like oblong, round, oval and reinform etc. Quantitative traits differed significantly and genotypes had variations in number of tubers, plant height and leaf.

Similarly, high genetic diversity was recorded for weight of tuber while low for germination, eyes of tubers and number of stem. Similarly, outcomes of biplot analysis displayed two different categories of genotypes on the bases of their qualitative and quantitative traits. These results are in lines with (Saint Pierre et al., 2008). Correlation analysis helps to determine the effective traits which are suitable and superior. Therefore, it helps the plant breeders regarding improvement in genetic traits particularly yield (Leilah & Al-Khateeb, 2005).

From these results, it is obvious that important variables such as tuber yield/plant, number of tubers/plant, weight of tubers, height of plant, sprouting of plants and leaves/plant might be taken during selection of parents during hybridization program and for development of elite lines (Mondal, Hossain, Rasul, & Uddin, 2007). In potato, total economic yield and its contributing traits are important and should be recommend as one of the best breeding strategy for genetic improvement of tuber yield in potato (Ahmadizadeh, Mostafa and Felenji, 2011). Similar results are indicated by several researchers like Haydar *et al.* (2007), as well as Lohani *et al.* (2012). Tairo *et al.* (2008) found the variability in Tanzanian landraces. Lohani *et al.* (2012) observed that first 11 components explained 96.25 % variation. The maximum variation of 18.78 % was explained by first latent vector followed by 16.34 % (second vector) and 13.30 % (third vector). Based on the component values, the location of genotypes and their grouping were determined in top of bi-plot (Figure 4). Therefore, according to bi-plot figures,23, 29, 31, 37, 38, 39, 41, 42, 43, 44, 45, 46, 47, 48, 49, 50, as well as 2, 5, 6, 7, 8, 9, 13, 15, 16, 17, 19, 20, 21, 23, 27, 28 and 30 identified as the best genotypes as these genotypes grouped in positive part of the bi-plot. Bi-plot had been used by many researchers such as (Ahmadizadeh, Mostafa and Felenji, 2011) in potato, (Afuape, Okocha, & Njoku, 2015) and Sivakumar *et al.* (2007) in sweet potato. It is essential to choose the genotypes, which have superior characters in genetic diversity and agronomic importance characters (Pandey, Singh, & Manivel.., 2005). Karaca (2004) selected 9 out of 63 genotypes in potato on the basis of maturity time, tuber shape and plant height. Thus, from the above investigation it can be concluded that principal component and biplot analysis in potato cultivars facilitated in identifying desirable traits and their relationship with yield and reliable classification of genotypes. The above variables might be taken into consideration for effective selection of parents during hybridization program. A good hybridization program can be initiated by the selection of cultivars from the bi-plot figure (Figure 4), with which we can identify core genotypes and relationship with morphological traits for specific breeding purposes.

## Conclusions

Considerable variability is present in the studied potato genotypes and these may be used start a good hybridization breeding programme by the selection of cultivars from the PC1 and PC2, genotypes and their relationship with morphological traits.

